# A method for the fast and photon-efficient analysis of time-domain fluorescence lifetime image data over large dynamic ranges

**DOI:** 10.1101/2021.08.11.455983

**Authors:** Romain F. Laine, Clemens F. Kaminski

## Abstract

Fluorescence lifetime imaging (FLIM) allows the quantification of subcellular processes *in situ*, in living cells. A number of approaches have been developed to extract the lifetime from time-domain FLIM data, but they are often limited in terms of dynamic range, speed, photon efficiency or precision. Here, we focus on one of the best performing methods in the field, the center-of-mass (CMM) method, that conveys advantages in terms of speed and photon efficiency over others. In this paper, however, we identify a loss of photon efficiency of CMM for short lifetimes when background noise is present. We subsequently present a new development and generalization of the CMM method that provides for the rapid and accurate extraction of fluorescence lifetime over a large lifetime dynamic range. We provide software tools to simulate, validate and analyze FLIM data sets and compare the performance of our approach against the standard CMM and the commonly employed least-square minimization (LSM) methods. Our method features a better photon efficiency than standard CMM and LSM and is robust in the presence of background noise. The algorithm is applicable to any time-domain FLIM dataset.

## Introduction

Fluorescence lifetime imaging (FLIM) provides a functional readout of phenomena occurring at the molecular level and, in contrast to intensity-based imaging, informs not only on the location of a fluorescent label but also its local environment. It is now widely used in biological research to quantify a plethora of cellular parameters, including ion concentrations, temperature or viscosity (1–4). FLIM has been implemented in numerous modalities, which include point-scanning and wide-field imaging methods (5) both in the time- and frequency-domains (6). In time-domain approaches, the lifetime is commonly extracted from FLIM data using non-linear least square minimization (LSM), for which a number of open-source packages are available (7, 8). Typically, LSM requires the acquisition of many temporal gates to yield accurate measurement of the lifetime, and this leads to long acquisition times. Some methods, such as rapid life-time determination (RLD), allow video rate FLIM (9) but are precise only over a comparatively small range of lifetimes (10). However, with recent technological advancement, in particular in the complementary metal-oxide-semiconductor (CMOS) and single-photon avalanche diode (SPAD) technologies, FLIM has gained a significant improvement in temporal resolution without loss in lifetime precision (11, 12), essentially through efficient parallelization in order to circumvent pile-up effects (13). These developments have renewed the interest in fast and reliable algorithms for lifetime determination. One such method is the so-called center-of-mass method (CMM), in which the first moment (center-of-mass) of the fluorescence decay is used as a lifetime estimator. The CMM algorithm is non-iterative, computationally efficient, and has been implemented on-chip, permitting on-the-fly lifetime estimation (14, 15).

In the CMM method, a temporal window (the analysis window) within the acquired decay time-range (the acquisition window) is chosen to compute the center-of-mass. We recall that the CMM method is based on the calculation of the first moment (center-of-mass, *CM*) of a fluorescence decay:

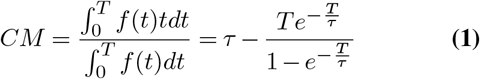

where 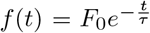 is the fluorescence decay, *τ* is the fluorescence lifetime and *T* the measurement window, over which the signal is measured. For *T* ≫ *τ*, *CM* ~ *τ* as the second term vanishes, but for short measurement windows, a correction needs to be applied to infer *τ* and take into account the finite size of the measurement window. This can be achieved using either an iterative approach or a look-up table (16).

In this paper we identify that, when background noise is present, lifetimes that are short with respect to the size of the analysis window are evaluated inefficiently, leading to the imprecise estimation of the lifetime. Exploiting the flexibility in the choice of analysis window, we present a generalization of the CMM method called F3-CMM to correct for this effect by applying the fusion of three lifetime images obtained with adapted temporal analysis windows. Our approach extends the dynamic range of CMM with high photon efficiency. The method potentially works for all time-domain FLIM datasets and could also be implemented on-chip, similarly to the standard CMM method (14).

## Results

### Photon efficiency of CMM and LSM lifetime estimations

To compare the performance of the CMM method with other commonly-used analysis methods, we modelled time-correlated single photon counting (TCSPC) data with Monte-Carlo simulations of photon arrival times (see Figure S1 and Materials and Methods for details). The software is distributed with this work and includes a Gaussian model of the instrument response function (IRF), photon noise, the effects of after-pulsing (17) and laser repetition rate. The after-pulsing (Ap) represents the fraction of photons in the background compared to those in the decay and is here a metric of the amount of noise on the background level. The software tool allows the simulation of realistic FLIM data that can be directly imported into our analysis software described further below, or into the commonly-used FLIMfit software (7). In a previous error analysis of the CMM method (14), simulations did not include the effect of background noise on algorithm performance, although background correction is an essential step in the CMM method (15). Here we use simulations for typical acquisition conditions for TCSPC (time window *T* =25 ns, 256 time bins, a Gaussian IRF centered at 3.2 ns with a standard deviation of 150 ps and containing a total of 108 photons) to estimate the quality of the lifetime estimation of F3-CMM and compare it to the common LSM method. A useful parameter to estimate the photon efficiency of a lifetime method is the F-value (18), which compares the signal-to-noise ratio (SNR) for a photon counting method to the precision of the measured lifetime:

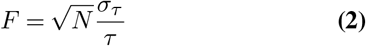

where *N* is the total number of photons, *τ* is the fluorescence lifetime and *σ_τ_* is the standard deviation of the lifetime measurement obtained from repeated measurements. For an ideal method, the photon efficiency reaches the minimum value of *F* =1, for other cases *F* > 1.

Figure 1 shows fluorescence lifetime images from simulations of a sample with uniform lifetime (2.5 ns). The simulation features a gradient of photon counts (from 100 to 5,000 photons) with background noise levels of Ap=0% and Ap=5% and was analysed with both standard CMM and LSM methods. Panel (c) of Figure 1 clearly shows the effect of photon noise on lifetime precision, introducing errors that increase from ~50 ps to ~300 ps as photon counts are reduced from 5,000 to 100. It shows, however, that no discernible bias is introduced by the CMM estimation across the signal range investigated.

**Fig. 1.**
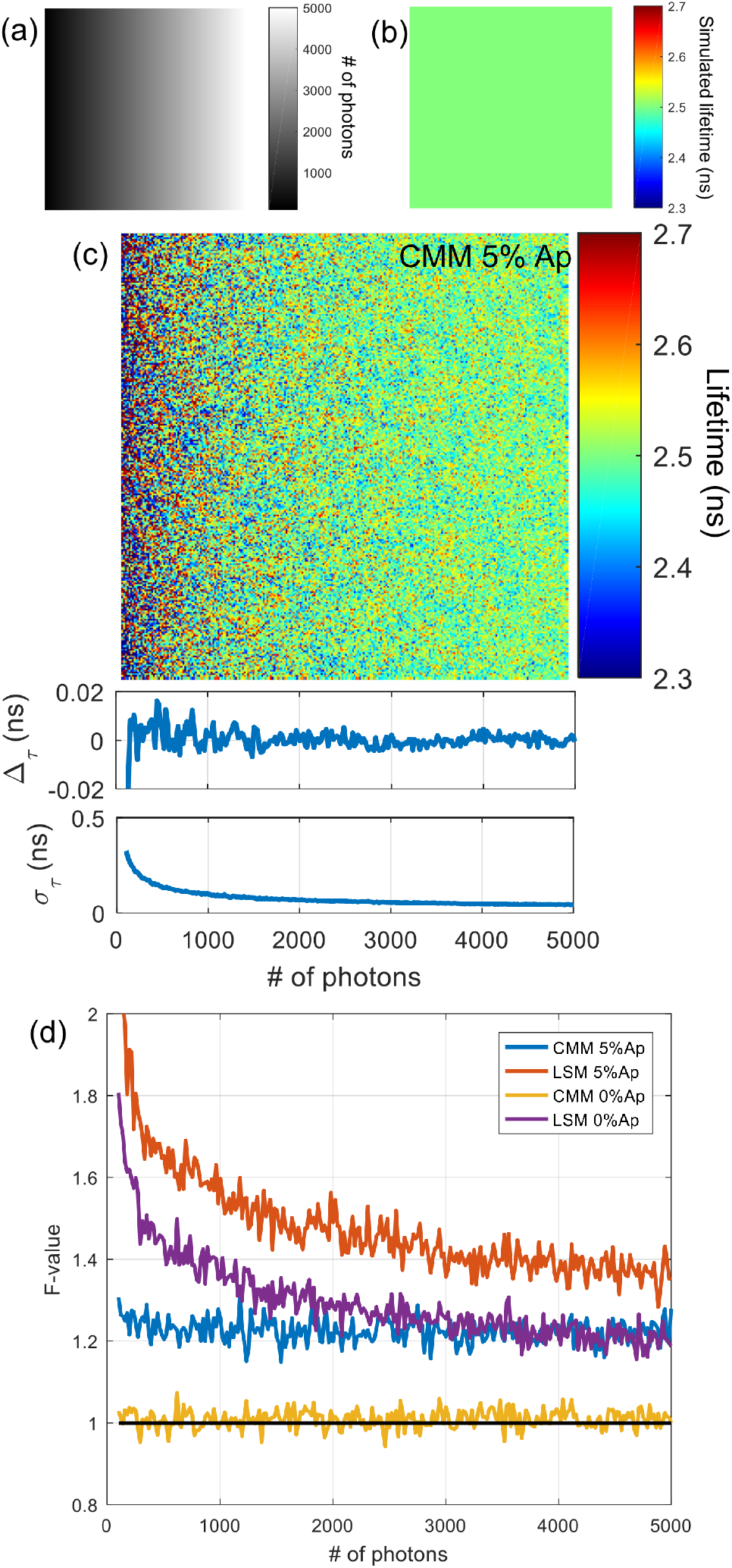
Effect of photon counts on lifetime determination in presence and absence of background noise. (a) Image of the total photon counts showing the gradient of photons from left (100 photons) to right (5,000 photons). (b) Uniform simulated lifetime image (2.5 ns). (c) Top: Lifetime image obtained from CMM analysis with 5% after-pulsing. Middle: Deviation of measured lifetime from the simulated lifetime (Δ_*τ*_) as a function of the number of photons (from 1,024 repeats). Bottom: Standard deviation of the measured lifetime (*σ_τ_*) as a function of the number of photons (from 1,024 repeats). (d) F-value as a function of the number of photons (from 1,024 repeats) for CMM and LSM in presence (5% Ap) and absence of background (0% Ap). When 5% after-pulsing is applied, the background is corrected by removing the average of the first ~20 times gates for CMM and by background level fitting for LSM. Ap: after-pulsing. The analysis window used the full acquisition window: 0-25 ns.

In Figure 1 panel (d), the F-value is plotted as a function of photon counts, for two levels of background noise and for both LSM and CMM methods. We observe that CMM determines the lifetime with constant photon efficiency across the whole range of counts. When background noise is present at Ap=5% the F-value increases from ~1 to ~1.23 from the case where no background noise is present. The LSM method, on the other hand, has lower photon efficiency which furthermore varies with photon counts. Figure S2 shows further data from these simulations, including lifetime images, accuracy and precision plots and data used to generate the F-value graphs in Figure 1(d).

The F-value does not take into account the potential presence of bias (loss of accuracy) as highlighted by Li *et al.* (14). We introduce an extension of the F-value in a format that takes account of both precision and accuracy of the method:

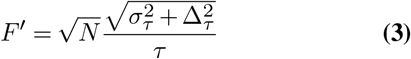

Here Δ*_τ_* is the difference between the mean of the measured and simulated lifetime. In combination, the F-value and F’-value can be used as figures of merit in simulated data to investigate the overall performance of a lifetime estimation method. We then investigated the photon efficiency of the methods as a function of the lifetime extracted. Figure 2 shows plots of both F and F’-values as a function of the simulated lifetime for the CMM and LSM methods in the presence and absence of background noise, at 5% and 0% after-pulsing, respectively. From Figure 2, we observe that the F’-value highlights a loss of accuracy of the LSM method at short lifetimes (< 500 ps). We also note that, in absence of background, the CMM method performs close to optimally (*F* ~ 1) for lifetimes ranging from 0.5 to 4 ns and better than the LSM method. We note that LSM suffers more strongly from the presence of background than CMM in the lifetime range > 2 ns. However, the opposite is true at shorter lifetimes (< 2 ns), where the precision of the CMM lifetime estimation rapidly worsens.

**Fig. 2.**
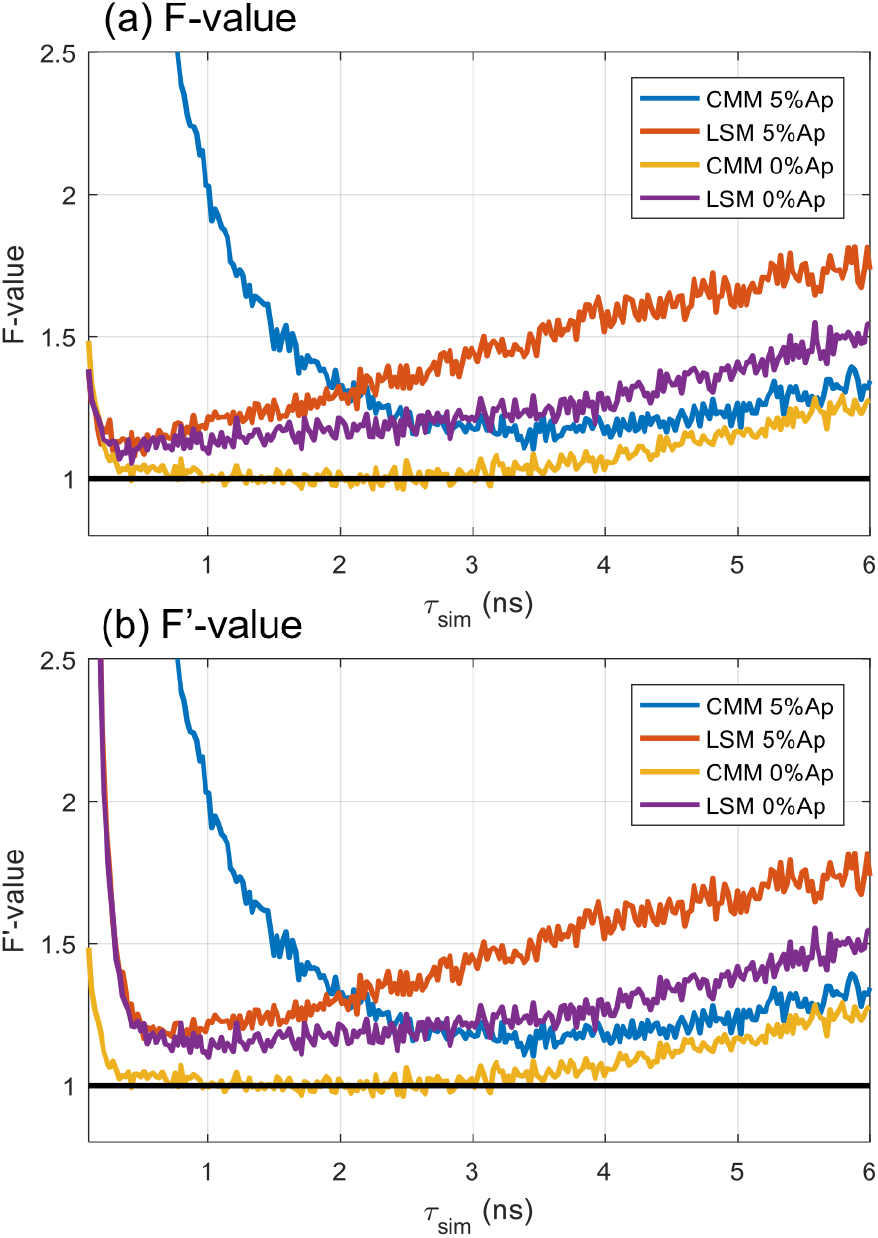
F-value (a) and F’-value (b) as a function of the simulated lifetime (*τsim*) for the CMM and LSM methods. The TCSPC decays were simulated with 5,000 photons with 5% or 0% after-pulsing background. When 5% after-pulsing is applied, the background is corrected by removing the average of the first ~20 times gates for CMM and by background level fitting for LSM. The F and F’ values were estimated from 1,024 repeats. The analysis window used the full acquisition window: 0-25 ns. Ap: after-pulsing.

Figure 2 shows, however, that the CMM method suffers from a loss of photon efficiency at short lifetimes (here < 2.5 ns). The reason for this is that, for short lifetimes, the background noise in the tail of the fluorescence decay accumulates and becomes a dominating source of error in the lifetime estimation.

### Improvement of CMM performance by adaptive windowing

The loss of photon efficiency of the CMM method in the short lifetime range can be mitigated through the use of shorter analysis windows, reducing cumulative noise and thus improving the precision of the method for short life-times. In Figure 3 (panel (a) and (b)), we show the F and F’-values for the CMM method for three different analysis windows (*T_a_* = 0-25 ns, 0-12.5 ns and 0-6.25 ns).

**Fig. 3.**
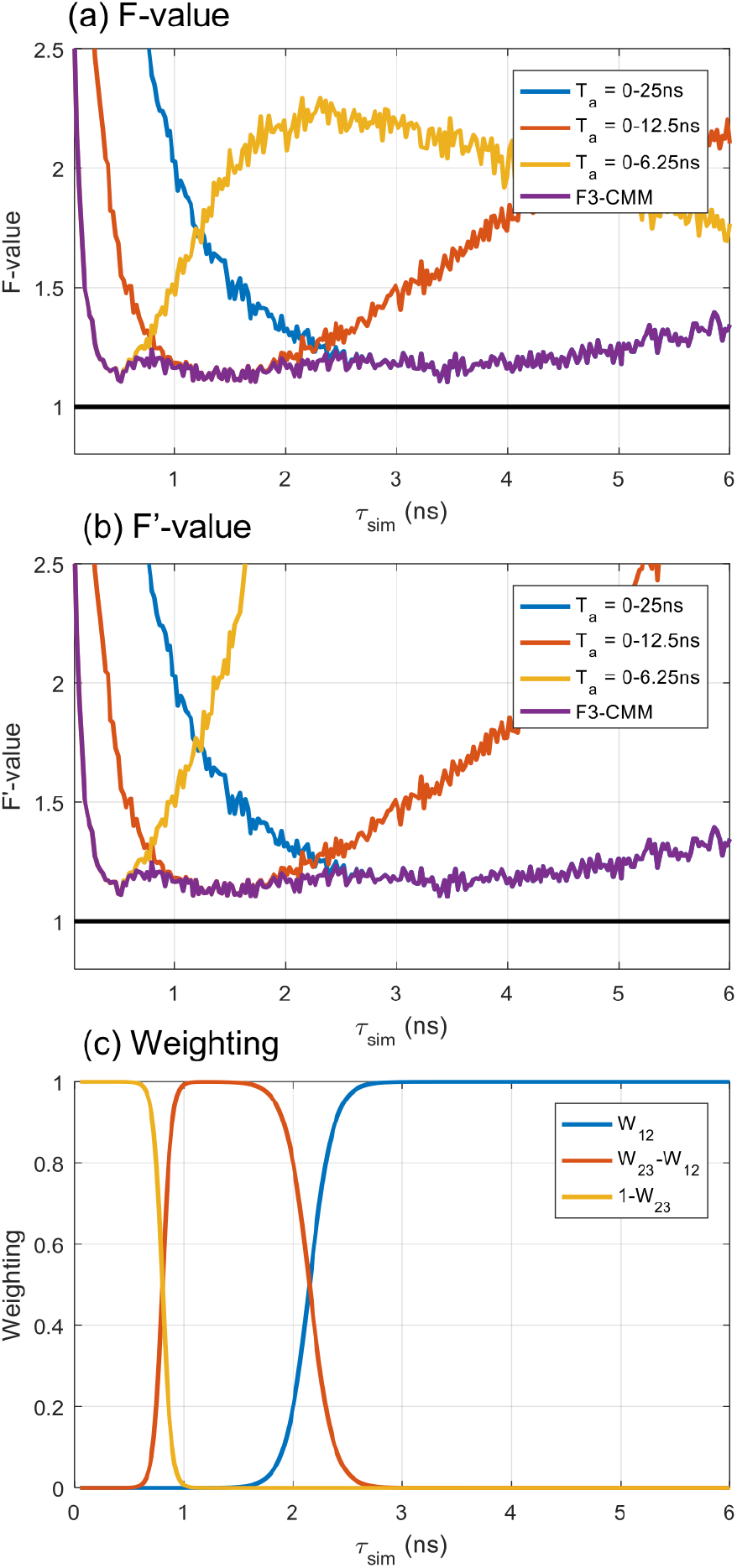
F-value (a), F’-value (b) and weighting factors used for F3-CMM (c) as a function of the simulated lifetime (*τ_sim_*). The CMM method was computed with 3 different analysis window sizes. The TCSPC decays were simulated with 5,000 photons with 5% after-pulsing background. The F and F’ values were estimated from 1,024 repeats. Ap: after-pulsing.

We observe that each analysis window performs optimally only for a certain range of lifetimes. However, no single analysis window performs well over the complete range of lifetimes considered here. This then highlights a problem with the CMM method since within a single FLIM acquisition lifetimes can vary greatly from pixel to pixel. When using a wide analysis window, a large standard deviation is obtained for short lifetimes. However, when using a small analysis window an inaccurate (biased) lifetime is obtained for long lifetimes. Here, to address this problem, we introduce the F3-CMM method, which combines the lifetime estimation from all three analysis windows and produces a composite lifetime image with improved precision and accuracy throughout the complete range of lifetime. The purple curve in Figure 3 was obtained from a weighted average of the results obtained from all 3 analysis window sizes following:

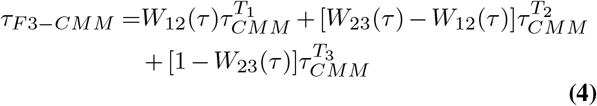

where *T*_1_, *T*_2_ and *T*_3_ correspond respectively to the analysis windows 25, 12.5 and 6.25 ns, *W_ij_* are the lifetime-dependent weighting factors, computed from

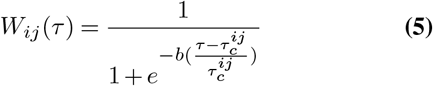

where 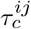 is the lifetime cut-off that separates analysis windows *T_i_* and *T_j_* and *b* is a ‘blending factor’ that determines the sharpness of the transition between adjacent weighting factors.

However, the weighting factors are lifetime-dependent and therefore cannot be directly computed in real data from a sample of unknown lifetime. We find that the optimal weighting factors can be well estimated from the CMM (*T* = 12.5 ns) as it is sufficiently accurate and precise in the region of the cut-off lifetimes (see Figure 3). Therefore, we use

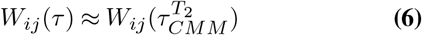

We note that an iterative method could be used to estimate the optimal weighting factors, instead of using the medium size window analysis but we do not expect this to lead to a significant improvement in the lifetime estimation.

In order to determine the lifetime range appropriate for a given analysis window size (and therefore the lifetime cut-off 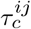), we plot the F’-value as a function of the ratio of the lifetime and the analysis window size as shown in Figure 4. This leads to a plot, the shape of which is invariant with analysis window size and which is well described by a rational function of the type

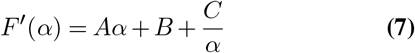

**Fig. 4.**
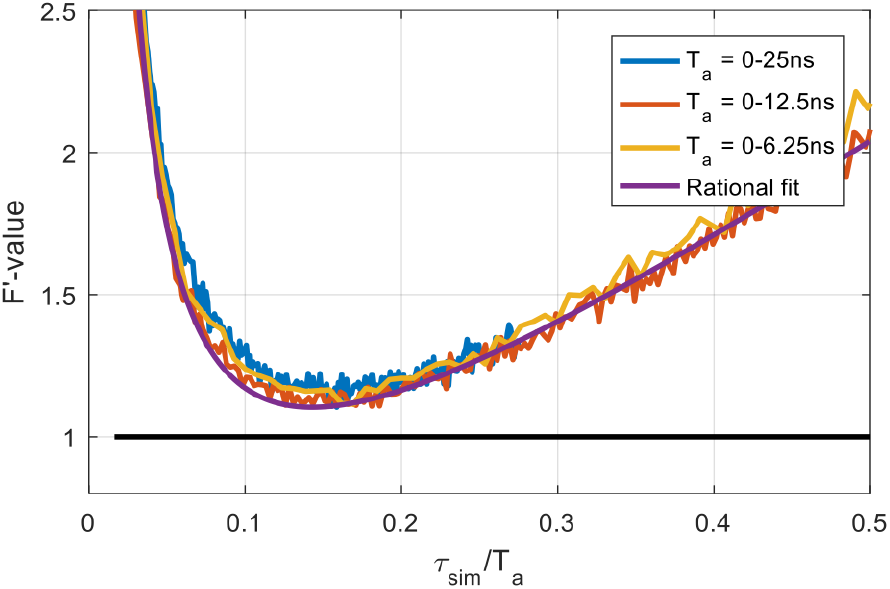
F’-value as a function of the ratio of the simulated lifetime and the analysis window with 3 analysis window sizes. The TCSPC decays were simulated with 5,000 photons with 5% after-pulsing background. The F and F’ values were estimated from 1,024 repeats. The rational fit is of the form: 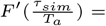 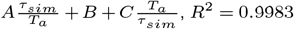, *R*^2^ = 0.9983.

This analytical description may be used to determine lifetime cut-offs 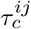 between two analyses windows *ij*, defined as the lifetime where the two curves cross and therefore

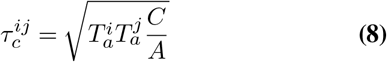

where 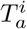 represents the effective analysis window size. The parameters *C* and *A* are relatively insensitive to the amount of after-pulsing (see Figure S3) and can therefore be estimated from the F’-value plot as a function of lifetime. For all results presented here, we used: A = 3.218 and C = 0.07339. We first tested the performance of our method *in silico*, by simulating TCSPC image data with a lifetime gradient. These simulated data were subsequently analyzed with F3-CMM and CMM methods. The results are shown in Figure 5. As expected, each of the 3 analysis window performs well (low noise and low bias) only in specific regions of the lifetime map. The large window (*T_a_* = 0-25 ns) shows large noise at low lifetimes (0-1 ns range) and the small analysis window (*T_a_* = 0-6.25 ns) has large bias towards shorter lifetime in the long lifetime region (> 2 ns). The F3-CMM method, on the other hand, is able to estimate the lifetime correctly over the entire lifetime range.

**Fig. 5.**
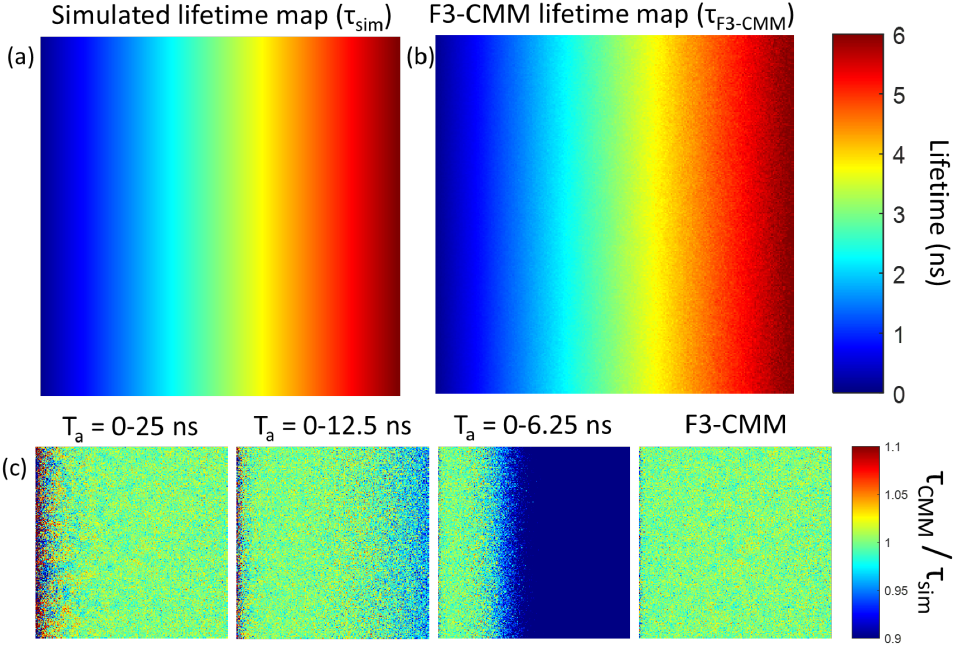
Performance analysis of the F3-CMM method for simulated data and comparison with the standard CMM method. (a) Simulated lifetime map (representing ground truth). (b) Lifetime map extracted by the F3-CMM method. (c) Comparison of CMM and F3-CMM results with ground truth, expressed as the recovered lifetime divided by the simulated lifetime. The first 3 panels show CMM data for different analysis windows. The 4th panel corresponds to F3-CMM. The TCSPC image data was simulated with 5,000 photons with 5% after-pulsing background.

We also generated *in silico* datasets to simulate challenging acquisition conditions. In Figure S4, we present a dataset containing 3 different lifetime values (0.5 ns, 1.5 ns and 3.5 ns) and variable signal photon numbers (~20, ~50 and ~120 respectively, corresponding to ~5 photons in the maximum bin in all decays) with a 5% after-pulsing background. It is clear that F3-CMM offers improved performance over CMM and LSM analyses in terms of precision and accuracy.

### Validation of the approach on experimental data

Next, we validated the method on experimental data. For this purpose, we used lifetime calibration solutions based on Rhodamine 6G dye solutions containing varying concentrations of potassium iodide (KI) as a fluorescence quencher (19). The calibration solutions obtained this way have been shown to provide a large range of lifetimes and allow titration of the quencher (6, 20). Here, we filled three transparent glass capillaries with varying mixtures of Rhodamine 6G and KI and imaged them side-by-side in a single field-of-view using our custom-built TCSPC confocal microscope (21). The results are shown in Figure 6. The recovered lifetime images and the histograms demonstrate the superior performance of the F3-CMM approach, leading to sharper lifetime distributions and less noisy images compared to LSM and standard CMM. The conventional CMM method leads to a broader distribution at short lifetimes (~0.3 ns), where there is also bias, evident from the large tail extending to lifetimes beyond 3 ns (see Figure 6(e)). This tail is absent in the F3-CMM approach.

**Fig. 6.**
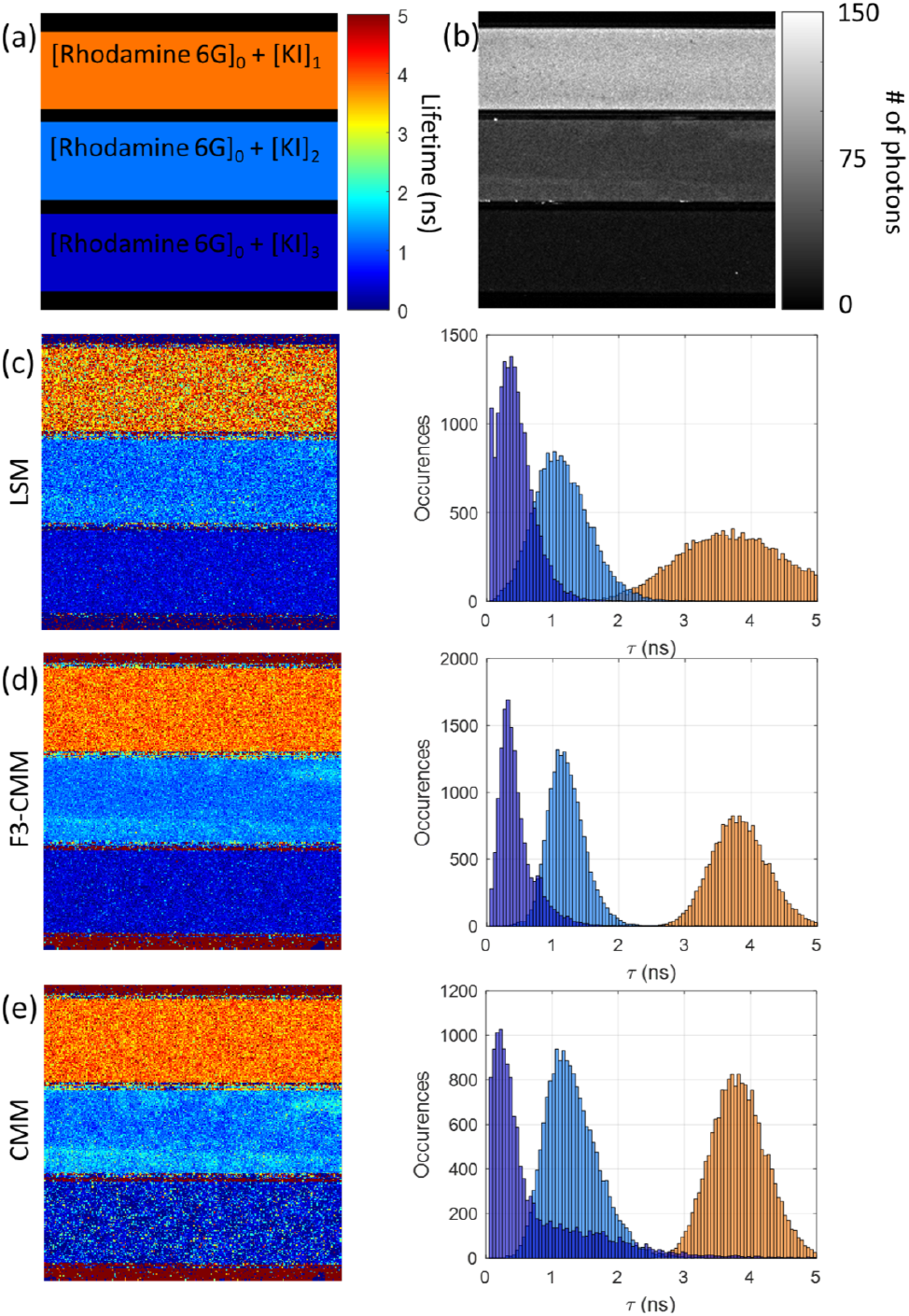
Comparison of F3-CMM, LSM and conventional CMM on Rhodamine 6G data. (a) Diagram representing the set-up of the capillary sample. Three rectangular capillaries filled with Rhodamine 6G and increasing concentrations of quencher KI were used. (b) Total intensity image obtained from the TCSPC dataset. (c-e) Recovered lifetime images (left) and corresponding lifetime histograms (right). The histograms correspond to a strip of 67 pixels wide centered on each capillary. The lifetime scale is indicated as in (a). The background in the images represented an after-pulsing in the range of 1-4%.

The means and standard deviations for the recovered life-times are compared in Table 1 for the different methods. Here again, the F3-CMM lifetime fraction map shows the best recovery of lifetime across the whole range of lifetimes. The lifetime values obtained for the conventional CMM, the LSM and F3-CMM methods are in good agreement with each other (~ 3.8, ~1.2 and ~0.3 ns respectively for capillary [*KI*]_1_, [*KI*]_2_ and [*KI*]_3_), but the standard deviations varied significantly. For the long lifetime, the standard deviation was halved by the CMM approaches compared to LSM. Also, CMM and F3-CMM performed identically, as expected since the conventional CMM method uses the full analysis window ideal for long lifetimes. For mid-range lifetime (~1.2 ns), the F3-CMM shows an improvement of 1.43-fold on the standard deviation over CMM. Finally, for the short lifetime (~0.3 ns), F3-CMM leads to a 2-fold improvement of the standard deviation over CMM. The estimated F-value shown in Table 1 quantitatively highlights the photon efficiency of the F3-CMM method.

**Table 1.**
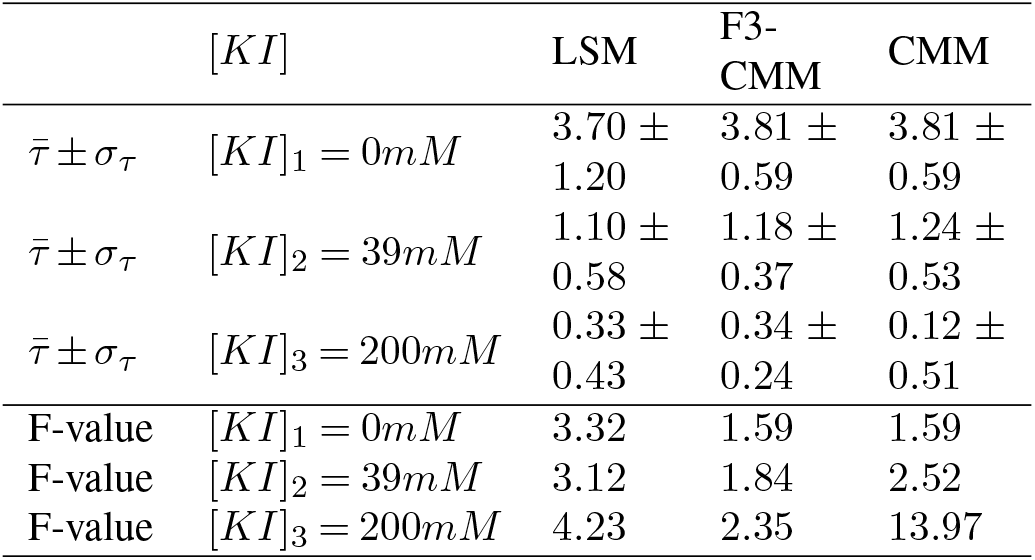
Table 1: Means and standard deviations of the lifetime estimated for each capillary for all three methods. The mean lifetime 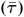 and its standard deviation (*σ_τ_*) were estimated by fitting a Gaussian function to the lifetime distribution. The F-value can then be estimated in each case by using the average number of photons in the decays from each capillary (105, 35 and 11 respectively for [*KI*]_1_, [*KI*]_2_ and [*KI*]_3_).

## Conclusions

Our simulations and experimental data clearly show the potential of CMM and, in particular, of the F3-CMM extension as a method to accurately and precisely estimate the fluorescence lifetime from single exponential decays over a wide lifetime dynamic range. The photon efficiency (indicated by the F’-value) is higher than for LSM for lifetime ranges typically encountered in biological fluorescence experiments, improving precision and accuracy of the lifetime determination for the same photon budget. Generally, the CMM algorithm is also faster than state-of-the-art LSM implementations and has been implemented ‘on-chip’ with FLIM instrumentation (14). The disadvantage of the CMM approach is that multiexponential decays cannot be distinguished and CMM will provide an estimate of an average lifetime instead. If this is desired, a global analysis and multi-exponential fitting with LSM are powerful alternatives. However, the average life-time is often sufficiently informative to reveal functional information and, with appropriate calibration, is quantitative for the measurement of absolute ligand concentrations (1). F3-CMM is robust with respect to changes in background noise but the parameters used to compute the weighting factors can be adjusted if exceptionally large amounts of background noise are present. F3-CMM will be especially beneficial when background noise is present in the dataset, for example when using SPAD arrays, which typically feature dark count rates that are typically higher than that encountered with photomultiplier tubes or with hybrid detectors (15). In conclusion, we have shown that using an adapted analysis window with CMM lifetime estimation leads to excellent photon efficiency with high precision and accuracy over a large range of lifetimes. We have introduced the F’-value as a measure of overall photon efficiency that takes into account both the precision and the accuracy of the method. It also helps identifying the range of lifetime that the method can extract with high fidelity. In addition to its high photon efficiency, the speed afforded by F3-CMM offers potential for quantitative dynamic, live cell measurement applications and for on-chip, video-rate implementation of FLIM analysis.

## Materials and methods

### Monte-Carlo simulation of TCSPC dataset

TCSPC data were generated using a Monte-Carlo simulations in MATLAB from the probability density functions of the photon excitation and emission times, which were represented by the impulse response function (IRF) model and the fluorescence decay model, respectively. The cumulative probability densities were calculated from the probability densities as shown in Figure S1. For the photon emission time we used

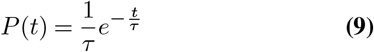

and

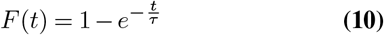

For the Gaussian IRF we used

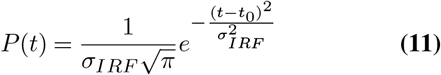

and

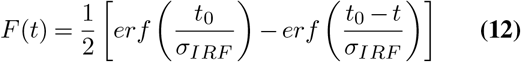

Two numbers *x*_1_(*i*) and *x*_2_(*i*) were generated by a pseudorandom number generator (PRNG) from a uniform distribution between 0 and 1 using the MATLAB function rand. These numbers were used to obtain the corresponding emission and excitation times, *t_em_* and *t_ex_* respectively, as shown in Figure S1. The photon arrival time is the sum of the two: *t_arr_ = t_em_ + t_ex_*. The number of photons in the background was determined from the after-pulsing (Ap) and the total number of photons (*N*): *N_b_ = ApbN*. The background noise level was simulated by randomly distributing (uniformly distributed across the acquisition window) the background photon arrival times across the acquisition window. The *N* + *N_b_* arrival times were binned to form a histogram of arrival times. Then, the arrival times beyond the acquisition window were wrapped around applying a modulo operation using the temporal period between laser pulses. The IRF was simulated by setting the emission time (*t_em_*) to zero. The simulation tool can generate a range of lifetimes and a range of photon number in an image of any size and then save it as 16-bit TIFF stacks using the OMERO MATLAB utilities. The stack can then be used for CMM analysis or imported into FLIMfit (7) for LSM analysis. The code is available on the author’s GitHub. https://github.com/Romain-Laine/TCSPC-image-simulation

### Computation of the CMM and F3-CMM algorithms

Prior to computing the center-of-mass, the backgrounds were removed from each decay by calculating the average of the first few time bins before the rising edge of the decay and subtracting it from the decay. Then, the lifetime extracted by the CMM method was estimated by calculating the centre-of-mass lifetime estimator, *τ_CM_*, of the corresponding decay with the chosen analysis window.

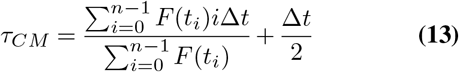

Where *F* (*t_i_*) is the fluorescence measurement in the temporal bin #*i* and Δ*t* is the bin size. The centre-of-mass of the IRF decay *τ_IRF_* was also computed in order to remove the effect of the IRF on the lifetime estimation. The CMM lifetime *τ_CMM_* was then obtained by subtracting the IRF lifetime.

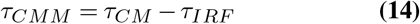

Then the lifetime corrected for the finite analysis window, 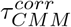, was computed by iterative method as follows.

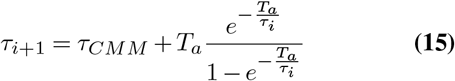

Where *T_a_* is the effective analysis window size and with *τ*_0_ = *τ_CMM_*. The effective analysis window is given by the difference between the total analysis window and *τ_IRF_*. In practice, we noticed that 10 iterations were sufficient to obtain unbiased estimates of the lifetime within the range of lifetimes considered here, therefore 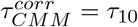. The F3-CMM is computed as described in the main text. A blending factor of *b*=20 was found to work well in most cases by providing a sharp transition between the lifetime images obtained from different analysis windows. The analysis was written as an open-source Fiji (22) macro tool and is distributed on the GitHub page of the author. https://github.com/Romain-Laine/F3-CMM-FLIM-analysis

### Rhodamine 6G FLIM imaging

Rhodamine 6G (Sigma, R4127) were prepared in increasing concentrations of potassium iodide (Sigma-Aldrich, 60400) in accordance to Hanley *et al.* (19). The dye and quencher mixture were freshly prepared and then used to fill up h ollow rectangle capillaries (CM Scientific, ID 0.10 × 1 .00mm). The capillaries were placed side by side on the microscope stage and imaged on a custom-made TCSPC microscope as previously described (21). An Olympus 2x 0.08 NA objective, 510 nm excitation wavelength and 560 nm emission wavelength (Semrock 560/25 filter) were used. The laser source was a Fianium SC400-4 set to a 20 MHz repetition rate. A 140s total integration time was used to measure the FLIM image.

## ACKNOWLEDGEMENTS

UK Engineering and Physical Sciences Research Council (EPSRC) (EP/L015889/1 and EP/H018301/1); Wellcome Trust (3-3249/Z/16/Z and 089703/Z/09/Z); UK Medical Research Council (MRC) (MR/K015850/1 and MR/K02292X/1); MedImmune; Infinitus (China) Ltd; UK Biotechnology and Biological Sciences Research Council (BBSRC) (BB/P027431/1).

The authors would like to thank Ian Munro and Sébastien Besson for providing the code for OME-tiff export and the OME team for maintaining the Bio-Format library. The authors would also like to thank Oliver Vanderpoorten and Súil Collins for suggesting using capillaries for the experimental datasets.

## Notes

https://github.com/Romain-Laine/F3-CMM-FLIM-analysis

https://github.com/Romain-Laine/TCSPC-image-simulation

### Competing Interest Statement

The authors have declared no competing interest.

