## Supplementary Information for "A method for the fast and photon-efficient analysis of time-domain fluorescence lifetime image data over large dynamic ranges"

Figure S1: Description of the TCSPC simulation.

Figure S2: Lifetime images, accuracy and precision plots for CMM and LSM as a function of photon counts.

Figure S3: F'-value as a function of the simulated lifetime in terms of fraction of the analysis window.

Figure S4: Comparison of CMM, F3-CMM with LSM on *in silico* data.

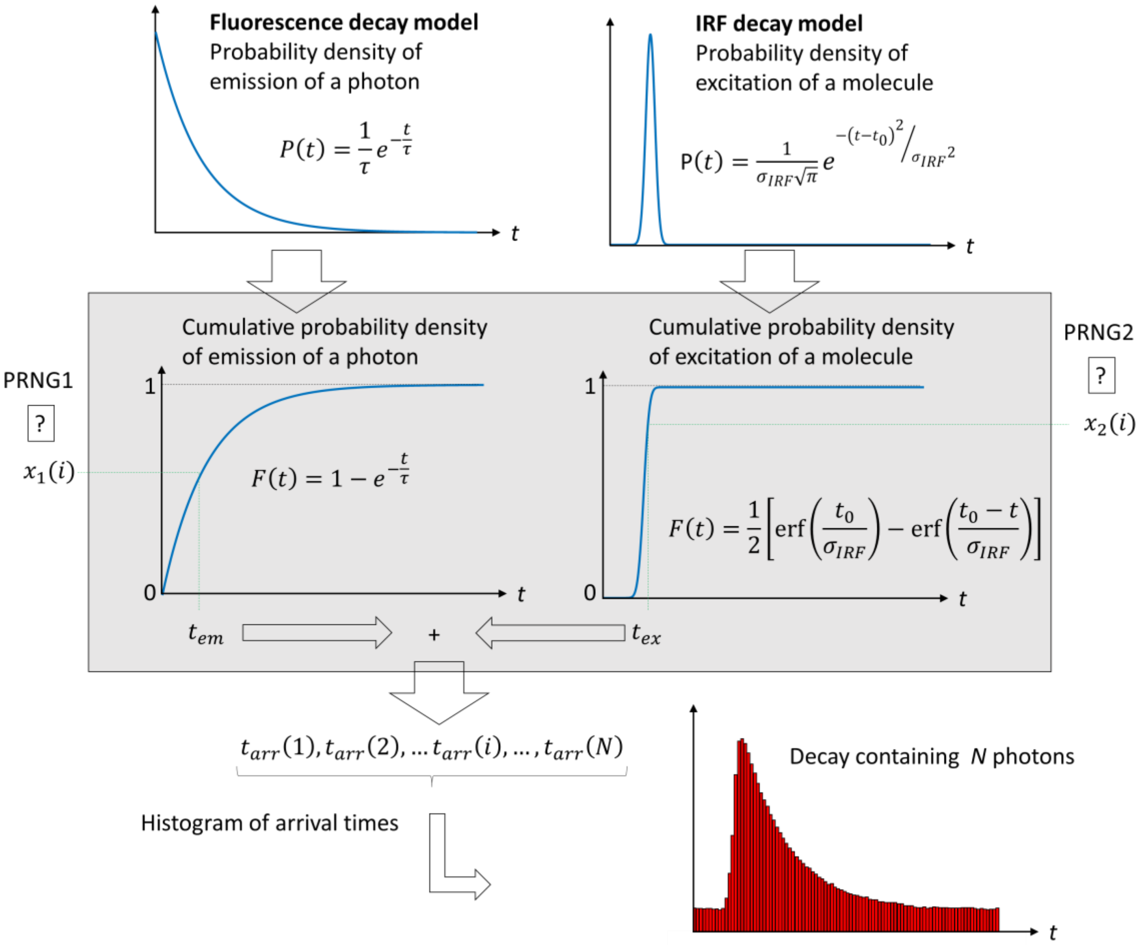


**Figure S1: TCSPC data simulation. PRNG: Pseudo-random number generator. N: total number of photons generated for the decay. $x_1(i)$ and $x_2(i)$ are the pseudo-random number generated for the calculation of the arrival time of the photon i.**

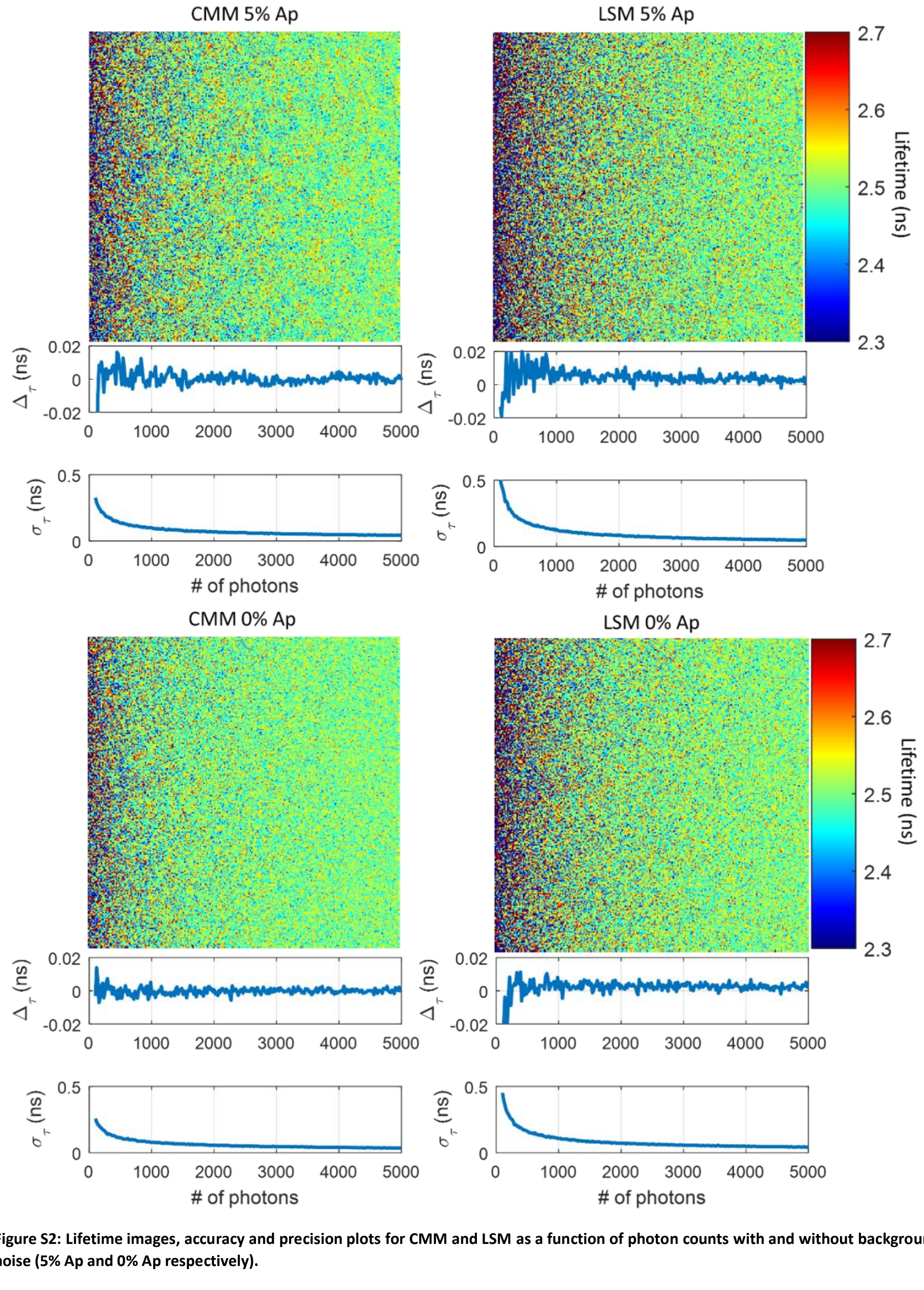


**Figure S2: Lifetime images, accuracy and precision plots for CMM and LSM as a function of photon counts with and without background noise (5% Ap and 0% Ap respectively).**

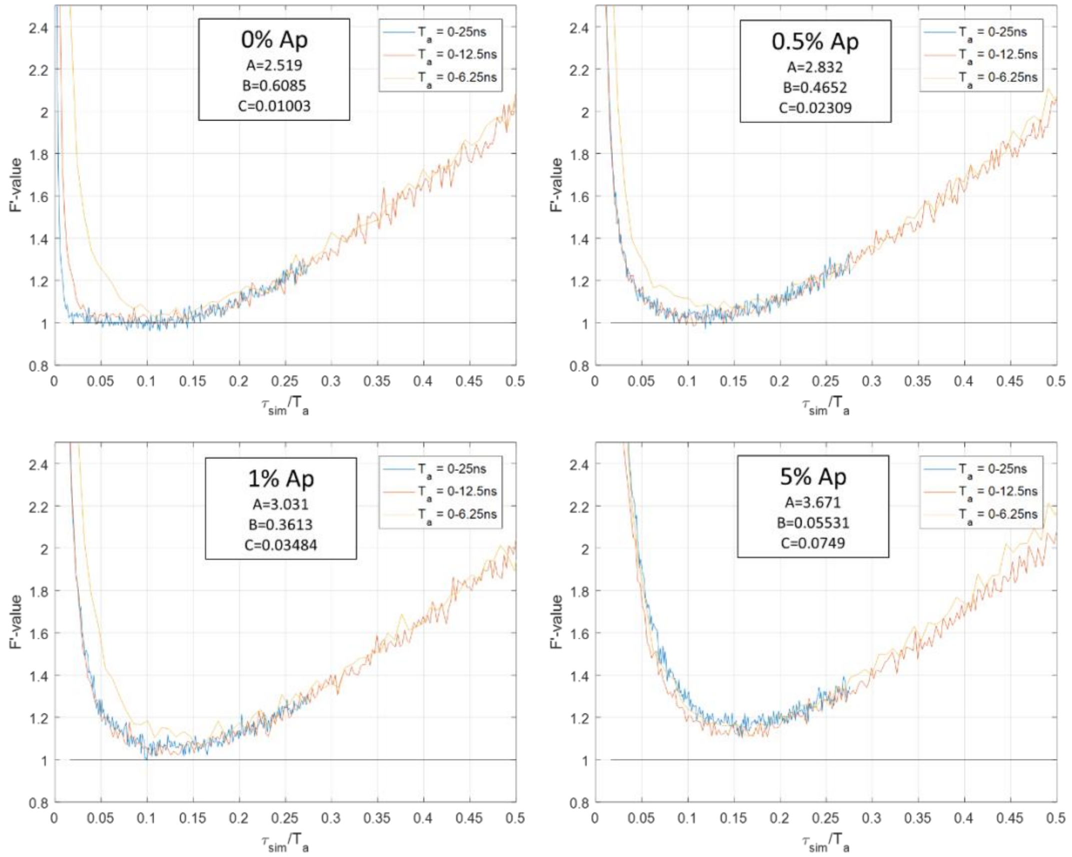


**Figure S3: F'-value as a function of the simulated lifetime in terms of fraction of the analysis window. Four different level of noise are investigated (0%, 0.5%, 1% and 5% of after-pulsing). N = 5,000 photons. The parameters of the rational function describing the curves are shown in the inset.** $F'\left({}^{\tau_{sim}}/_{T_a}\right) = A\,{}^{\tau_{sim}}/_{T_a} + B + C\,{}^{T_a}/_{\tau_{sim}}$

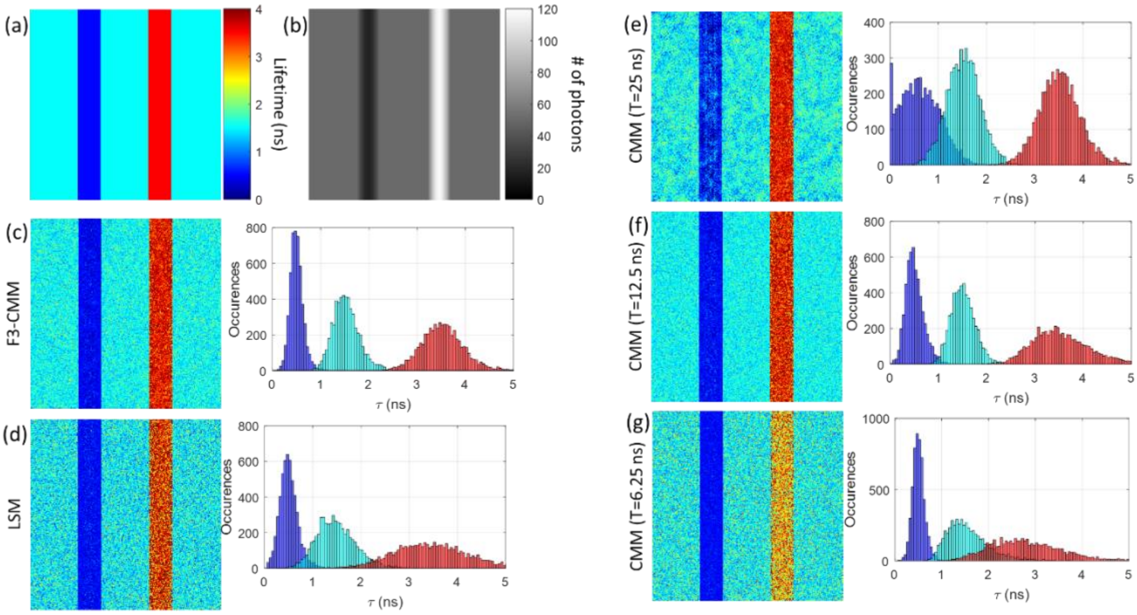


**Figure S4: Comparison of CMM, F3-CMM with LSM on *in-silico* data. The lifetimes were 1.5 ns, 0.5 ns and 3.5 ns. The number of photons were adjusted so that shorter lifetime had fewer photons, equalling to similar number of photons in the maximum bins for all lifetimes (~5 photons). The histograms of lifetime shown on the right of each lifetime image correspond to a strip of 20 pixels wide centred on the 0.5 ns strip (dark blue histogram), centred on the 1.5 ns strip (light blue histogram) and centred on the 3.5 ns strip (red histogram).**
